# RNAs as proximity labeling media for identifying nuclear speckle positions relative to the genome

**DOI:** 10.1101/300483

**Authors:** Weizhong Chen, Zhangming Yan, Simin Li, Norman Huang, Xuerui Huang, Jin Zhang, Sheng Zhong

## Abstract

Nuclear speckles are interchromatin structures enriched in RNA splicing factors. Determining their relative positions with respect to the folded nuclear genome could provide critical information on co-and post-transcriptional regulation of gene expression. However, it remains challenging to identify which parts of the nuclear genome are in proximity to nuclear speckles, due to physical separation between nuclear speckle cores and chromatin. We hypothesized that noncoding RNAs including small nuclear RNAs, 7SK and Malat1, which accumulate at the periphery of nuclear speckles (nsaRNA, <u>n</u>uclear <u>s</u>peckle <u>a</u>ssociated RNA), may extend to sufficient proximity to the nuclear genome. Leveraging a transcriptome-genome interaction assay (MARGI), we identified nsaRNA-interacting genomic sequences, which exhibited clustering patterns (nsaPeaks) in the genome, suggesting existence of relatively stable interaction sites for nsaRNAs in nuclear genome. Posttranscriptional pre-mRNAs, which are known to be clustered to nuclear speckles, exhibited proximity to nsaPeaks but rarely to other genomic regions. Furthermore, CDK9 proteins that localize to the vicinity of nuclear speckles produced ChIP-seq peaks that overlapped with nsaPeaks. Our combined DNA FISH and immunofluorescence analysis in 182 single cells revealed a 3-fold increase in odds for nuclear speckles to localize near an nsaPeak than its neighboring genomic sequence. These data suggest a model that nsaRNAs locate in sufficient proximity to nuclear genome and leave identifiable genomic footprints, thus revealing the parts of genome proximal to nuclear speckles.

## Introduction

It is increasingly evident that positioning and organization of various subnuclear structures are critical for regulating gene expression and therefore resolving spatial organization of nuclear components has become a central task to nucleome research ^1^. Nuclear bodies, previously known as interchromatin structures, typically exhibit non-overlapping spatial distributions with the genome ^2,3^. With an exception of nucleoli which are positioned near ribosomal DNA ^4^, it remains challenging to identify the genomic sequences near most of the nuclear bodies, especially nuclear speckles ^5^. Chromatin immunoprecipitation sequencing (ChIP-seq) targeting nuclear speckle core proteins rarely produces reproducible peaks ^6^, likely due to lack of stable physical interactions between nuclear speckle core proteins and chromatin ^6–8^.

Advanced imaging technologies including super-resolution imaging have started to reveal the multilayer structure of nuclear speckles, with proteins SC35 and SON at the center ^9^ and nuclear speckle-associated noncoding RNAs (nsaRNA) including snRNA and Malat1 at the peripheral ^9^, as well as pre-cursor mRNAs (pre-mRNA) accumulated at the peripheral ^10,11^. In addition, distribution of Cdk9-cyclin T1 complex correlates with nuclear speckles ^12,13^ but more often extends beyond the periphery of nuclear speckles ^7^ (Figure 1A). A number of other proteins are associated with nuclear speckles ^7^, however it remains unclear whether their distribution corresponds to specific layers. The microscopic observation that noncoding RNAs located to the outer layer of nuclear speckles ^9^ led us to hypothesize that these peripheral noncoding RNAs may be present in sufficient proximity to nuclear genome, leaving identifiable proximal sequences as their genomic footprints. Hereafter, we call this hypothesis the “nsaRNA proximity” hypothesis.

**Figure 1.**
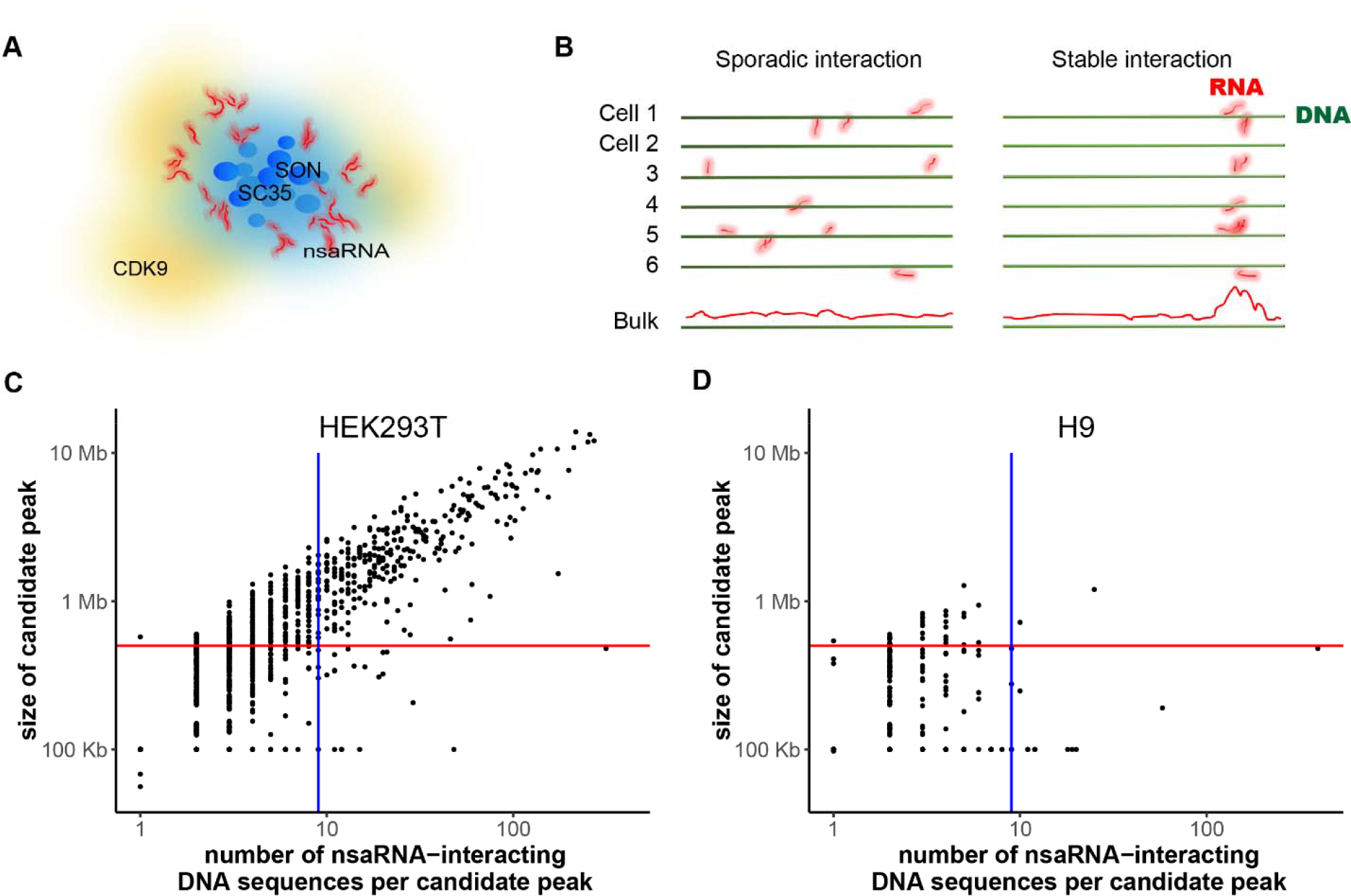
DNA interaction sites of nsaRNAs. (A) A cartoon of multilayer structure of nuclear speckles. (B) Models of RNA-chromatin interaction in single cells, including sporadic interaction model and stable interaction model. (C-D) Candidate peaks of nsaRNA-interacting DNA sequences in the genome. The number of nsaRNA-interacting DNA sequences (x axis) is plotted against cluster size (y axis) for every candidate peak in HEK293T (C) and H9 hES cells (D). Vertical line: 9 reads. Horizontal line: 500 Kb.

The recent technology on global <u>ma</u>pping of <u>R</u>NA-<u>g</u>enome interactions (MARGI) enabled identification of interacting genomic sequences of chromatin-interacting RNAs ^14^. After crosslinking and genome fragmentation, MARGI ligates RNA, a linker sequence, and proximal DNA to form a RNA-linker-DNA chimeric sequence, which is subsequently converted to double stranded DNA and subjected to paired-end sequencing (see Figure 1 of ^14^). Because MARGI simultaneously assayed thousands of noncoding RNAs including nsaRNAs, we will leverage MARGI data to test the nsaRNA proximity hypothesis.

Resolving spatial organization of nuclear components requires connecting information through different length scales and data types. Microscopic analyses have revealed non-uniform three-dimensional (3D) distribution of several types of RNAs in the nucleus. Prominent examples include Xist RNA cloud in adult female cells ^15^, accumulation of ribosomal RNA (rRNA) in nucleoli ^16^, accumulation of small nuclear RNAs (snRNAs), Malat1 and pre-mRNAs in nuclear speckles ^10,11,17–21^. However, it remains a challenge to connect these microscopic findings with the latest information on 3D genome organization derived from genomics assays ^1^. This challenge lies partially in the different length scales that vary in orders of magnitudes. For instance, the protein core of a nuclear speckle varies from one to several micrometers in diameter ^7^ that is approximately 20% to 50% of the spread of metaphase chromosomes ^22,23^ or the diameters of chromosome territories ^23,24^. These relative sizes suggest genomic regions in proximity to nuclear speckles may be significantly larger than the typical sizes of ChIP-seq or ATAC-seq peaks. Nevertheless, the enrichment of Xist RNA on X chromosome revealed by imaging ^15^ was successfully corroborated by genomics technologies including RAP-seq ^25^ and MARGI ^14^, offering an example of convergent findings from imaging and genomics approaches. In this work, we tested our “nsaRNA proximity” hypothesis by combining microscopic information and genomics data, and aimed for establishing an RNA-based approach for identifying relative positions of the folded genome and subnuclear structures.

## Results

### MARGI captures proximity of nuclear rRNA to ribosomal DNA

We used the co-localization of nuclear rRNA and ribosomal DNA (rDNA, human ribosomal DNA complete repeating unit) in nucleoli ^16^ as a testbed system to verify the assumption that RNA-DNA ligation sequencing (MARGI) data reflects spatial co-localization of a group of nuclear body-associated RNAs with specific genomic sequences. We reanalyzed MARGI datasets from human embryonic kidney (HEK) cells (GEO: GSM2427902 and GSM2427903) and human embryonic stem (hES) cells (GEO: GSM2427895 and GSM2427896) ^14^, which yielded approximately 9.9 million and 5.6 million RNA-DNA sequence pairs, respectively (Table S1). To test whether rRNAs are enriched in the proximity of rDNA, we categorized the RNA-DNA sequence pairs by the RNA type (rRNA or other types) and by the DNA (rDNA or the rest of the genome (hg38)) (Table S1). Compared to other types of RNA, rRNA exhibited more than 400-fold increase of odds to ligate with rDNA in HEK cells (odds ratio = 404, p-value <10^−16^) and more than 1800-fold increase of odds in hES cells (odds ratio = 1,810, p-value <10^−16^), confirming that MARGI data reflected co-localization of nucleolus-associated RNA and DNA.

### nsaRNA-DNA interaction is cell type specific

We asked which genomic regions are in proximity to nsaRNAs. At single cell level, there are three possible answers (models) to this question, which are 1) lack of nsaRNA expression; 2) nsaRNAs do not stably locate in proximity of any specific genomic region in a single cell; 3) nsaRNAs are proximal to different DNA sequences in different single cells, however none of these DNA sequences are shared by the majority of the cells; 4) nsaRNAs are proximal to some DNA sequences and at least a fraction of these DNA sequences are shared by the majority of cells. Experiments with bulk cells could potentially differentiate the fourth model (stable interaction model) from its opposite (the first two models, collectively called sporadic interaction model) (Figure 1B) but cannot further differentiate the first three models. Under the sporadic interaction model, bulk cell analysis (MARGI) is not expected to identify nsaRNA-DNA interactions (Bulk lane, Figure 1B).

We used MARGI datasets to test the competing models. We reprocessed MARGI datasets generated from HEK and hES cells using the MARGI analysis pipeline (http://systemsbio.ucsd.edu/margi/) ^26^. This pipeline obtains the RNA-DNA read pairs with both ends uniquely mapped to the genome (hg38) and subsequently removes the “proximal” read pairs where the two ends were mapped to genomic locations within 2,000bp to each other. HEK and hES cells yielded 559,873 and 211,487 uniquely mapped RNA-DNA read pairs, respectively. In HEK cells, 14,941 pairs (2.5%) were nsaRNA-DNA pairs with the RNA end uniquely mapped to nsaRNAs (U1, U2, U4, U4atac, U5, U6, U6atac, U11, U12 ^27–29 30,31^, 7SK ^17,32^, Malat1 ^19^). In comparison, there were 1,867 pairs nsaRNA-DNA pairs in hES cells, corresponding to only 0.88% RNA-DNA pairs in hES cells. Compared to HEK, hES-derived read pairs exhibited 3 fold reduction in odds of being nsaRNA-DNA repairs (odds ratio = 3.1, Chi-squared p-value < 10^−16^), which is reminiscent of lack of nuclear speckle formation in hES cells where SC35 proteins and nsaRNAs are diffusely distributed in the nuclei ^33^.

In HEK, the nsaRNA-interacting DNA formed candidate peaks (Figure 1C, red curve in Figure 2). Analysis with Homer (v4.8.3) yielded a total of 295 broad peaks (nsaPeaks, Figure 5), which contained 10,771 (72%) of the nsaRNA-interacting DNA sequences (permutation p-value < 0.001). The sizes of nsaPeaks ranged from 100 kb to 13 Mb, on the same scale of nuclear lamina-associated domains (LADs) that span 10 kb to10 Mb ^34^. The clustering of nsaRNA-interacting DNA sequences in the genome is consistent with the stable interaction model. In hES, nsaRNA-interacting DNA sequences barely exhibited any clustering formation in the genome and yielded 2 broad peaks by Homer analysis. Adjusting for the total amount of candidate peaks and isolated nsaRNA-interacting DNA in each cell type (Figure 1D), hES cells exhibited more than 80-fold reduction in producing nsaRNA-interaction peaks as compared to HEK cells (odds ratio = 88.9, p-value < 10^−16^). The sporadic distribution of nsaRNA-interacting DNA sequences in hES cells is also consistent to the lack of SC35 clusters in hES cells (Sporadic interaction, Figure 1B). On the other hand, the 295 nsaRNA-associated broad peaks (nsaPeaks) identified in HEK cells exhibited the expected characteristics of the genomic regions close to nuclear speckles. We proceeded to test these genomic regions with two other types of nuclear speckle associated molecules, namely pre-mRNAs and CDK9 proteins.

**Figure 2.**
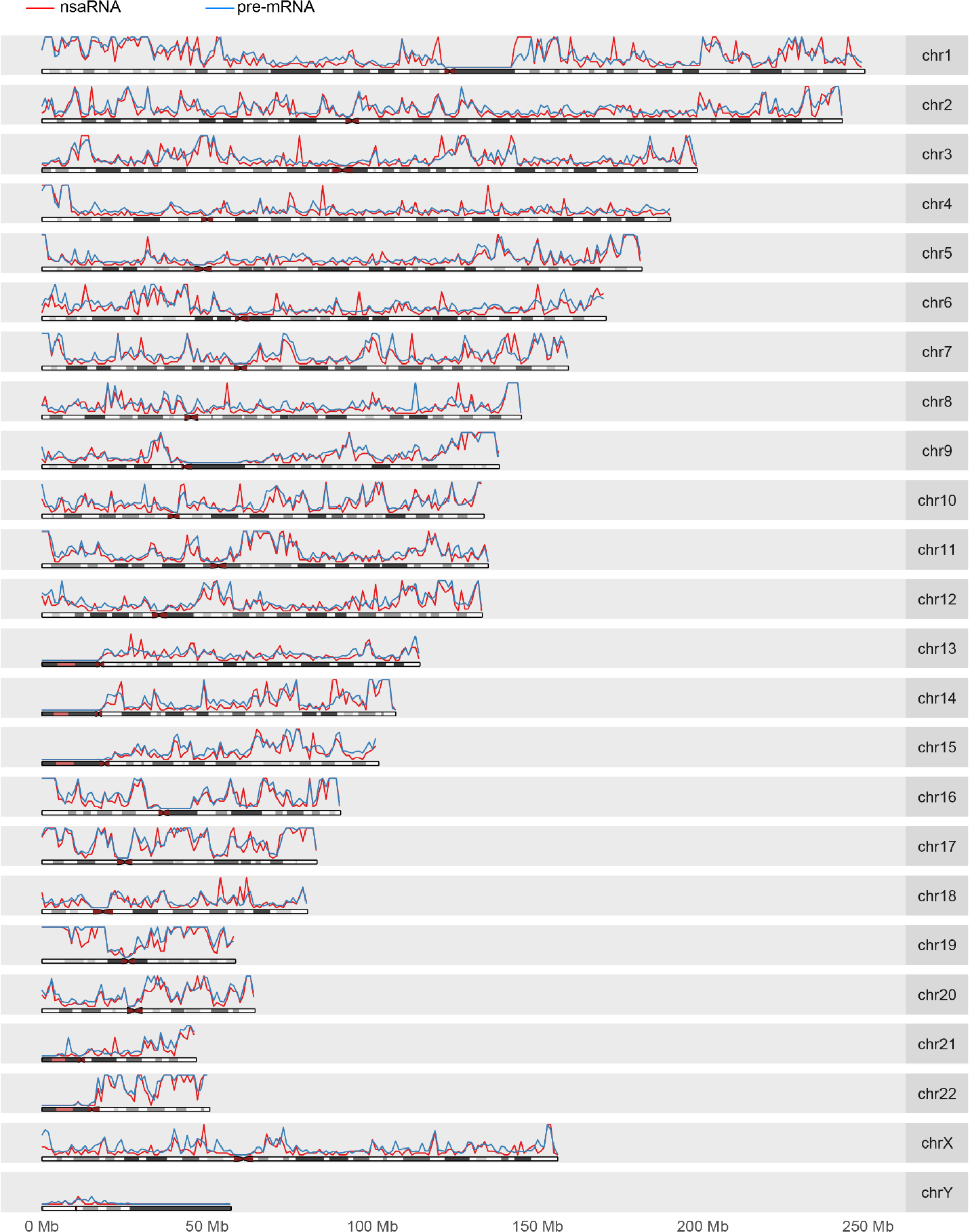
Genome-wide density distributions of nsaRNA-interacting DNA sequences (red curve) and pre-mRNA proximal DNA (blue curve).

### Pre-mRNA proximal regions overlap with nsaRNA-interacting DNA

If nsaPeaks are near nuclear speckles, other nuclear speckle associated molecules besides nsaRNAs may also exhibit enrichment in spatial proximity of nsaPeaks. Clustering of pre-mRNAs at nuclear speckle domains ^10,11,21^ offers another characteristic of nuclear speckles for testing nsaPeaks as the part of genome proximal to nuclear speckles. The key assumption of this test is that clustering of RNAs in 3D predicts clustering of their interacting genomic sequences in the genome. To test this assumption, we examined whether pre-mRNA-interacting genomic sequences exhibit clustering patterns or are sporadically distributed in the genome. We processed MARGI data from HEK cells ^14^ to identify pre-mRNA—DNA interactions. After removing the sequence pairs were likely produced from a nascent transcript and its own gene (the mapped RNA end and DNA end were within 2,000bp in the genome), 187,724 uniquely mappable sequence pairs representing pre-mRNA—DNA interactions were obtained. The 187,724 DNA ends of these sequence pairs were not uniformly distributed in the genome (blue curve, Figure 2), instead they concentrated to certain genomic regions, yielding 284 broad peaks (Homer v4.8.3, broad peak option) (p-value < 0.001, permutation test) (Figure S1). Taken together, pre-mRNA interacts with chromatin expect for its own genic location, and even more, pre-mRNA interacting DNA sequences are clustered in the genome, which corroborates with the idea that pre-mRNAs are clustered rather than diffusively distributed in the nucleus ^10,11,21^.

We compared nsaRNA-interacting DNA and pre-mRNA proximal DNA by genome-wide density distributions, broad peaks and genomic windows. The genome-wide distribution of pre-mRNA proximal DNA sequences exhibited remarkable similarity to the distribution of nsaRNA-interacting DNA (Figure 2). A total of 170 (57.6%) nsaPeaks overlapped with pre-mRNA broad peaks (Figures S1, S2A) (p-value < 0.001, permutation test). Finally, we broke the genome into equal-sized windows and calculated the densities of nsaRNA-interacting DNA and pre-mRNA proximal sequences in each window. These two densities profiles exhibited a genome-wide correlation (Spearman correlation = 0.957, p-value < 10^−16^) (Figure S2B-C). Taken together, pre-mRNA proximal genomic regions exhibited significant overlap with nsaRNA-interacting DNA, supporting the idea that nsaPeaks reflect the parts of genome near nuclear speckles.

### Correspondence of genome-wide binding profile of CDK9 and genome-wide distribution of nsaRNA-interacting DNA

We compared genome-wide binding profile of CDK9 to genome-wide distribution of nsaRNA-interacting DNA sequences. ChIP-seq of nuclear speckle core proteins has been regarded a questionable approach for identifying the relative positions of nuclear speckles and the genome ^1,8^, due to physical separation of nuclear speckle cores from chromatin ^7^. For example, suppose 95% of copies of a core protein, for instance SC35, were located at the nuclear speckle cores and the other 5% were sporadially distributed some of which are attached to chromatin, ChIP would select for the few chromain-associated SC35 rather than those at the nuclear speckle cores. To alleviate this documented concern, we resorted to CDK9 proteins that are distributed relatively broadly throughout the core and periphery of nuclear speckles ^7,12,13^ for a ChIP-seq analysis. And even so we did not anticipate many overlaps between CDK9 ChIP-seq peaks and nsaRNA-interacting DNA sequences. We identified a total of 6,517 CDK9 peaks from HEK293T cells (GEO: GSM1249897) ^35^ (MACS2) ^36^, of which only 551 (8.5%) located within 200bp of a nsaRNA-interacting DNA sequence. This overlap was statistically significant (p-value < 0.001, permutation test), consistent with the idea that CDK9’s distribution overlaps with nuclear speckles. However, the relatively small number of actual overlaps is reminiscent of the recognized challenge of using ChIP to identify nuclear speckle interacting genomic regains ^1,8^.

Considering that the 3D distribution of CDK9 is centered at nuclear speckles ^7,12,13^, we tested the possibility that CDK9 ChIP-seq peaks cluster to the same genomic regions as nsaPeaks. Indeed, genome-wide density distribution of CDK9 peaks (green curve, Figure 3) resembled the density distribution of nsaRNA-interacting DNA (red curve, Figure 3). In a control comparison, genome-wide density distributions of H3K9me3 (Encode Accession ID: ENCFF002AAZ) ^37^ and nsaRNA-interacting DNA exhibited a poor correlation (Pearson correlation = 0.03, Spearman correlation = 0.27) (Figure S3). To test whether CDK9 binding sites cluster to the same genomic regions as nsaPeaks, we identified a total of 262 CDK9 broad peaks (sizes range from 514,083 bp to 6,262,520 bp, median size = 1,328,930 bp) (Homer, v4.8.3) ^38^, of which 206 (78.6%) overlapped with nsaPeaks (p-value < 0.001, permutation test) (Figures S4A, S5). Next, we split the genome (hg38) into 3.08 million 1,000-bp windows, of which 0.44 million windows overlapped with CDK9 broad peaks, of which 0.32 million windows also overlapped with nsaPeaks, suggesting strong association (odds ratio = 11.5, p-value <10^−16^, Fisher’s exact test) (Figure S4B). Taken together, although CDK9 does not frequently bind to the exact sequences as nsaRNA-interacting DNA, CDK9 binding sites accumulated to nsaPeaks, corroborating with the idea that nsaPeaks reflect the portion of genome closer to nuclear speckles.

**Figure 3.**
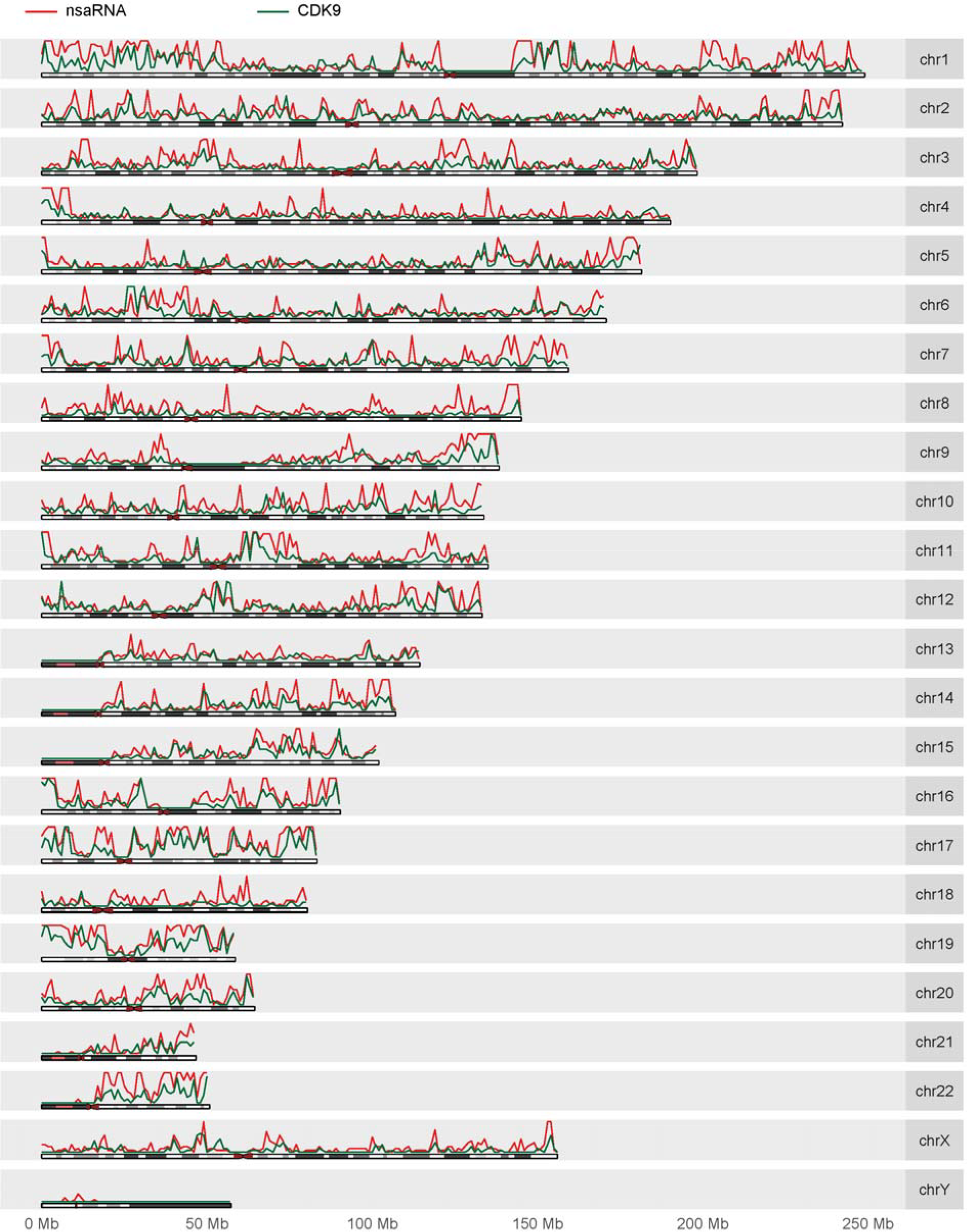
Genome-wide density distributions of nsaRNA-interacting DNA sequences (red curve) and CDK9 ChIP-seq sequences (green curve).

### Co-localization of SC35 clusters and nsaPeaks in single cells

We examined the proximity of nuclear speckles to nsaPeaks at single cell resolution using a combination of immunofluorescence staining of a nuclear speckle core protein SC35 and DNA fluorescent *in situ* hybridization (FISH) ^39^. We opted to use commercially validated FISH probes and we wanted the probes to be on the same chromosome arm. We identified a pair of probes satisfying these criteria on Chromosome 11 with one probe (BAC plasmid DNA) inside an nsaPeak (Empire Genomics: RP11-772K10, hereafter called nsaPeak probe) and the other probe outside nsaPeaks (Empire Genomics: RP11-908J16, hereafter called non-nsaPeak probe) (Figure 3A). We imaged 82 and 100 single cells with nsaPeak probe and non-nsaPeak probe, respectively. Each cell exhibited 1 to 3 FISH spots, consistent with pseudotriploidy of HEK293T cells, and 20 to 35 SC35 clusters (Figure 3B).

To minimize sensitivity of results to image analysis methods, we carried out two sets of analyses based on different analysis methods. First, we identified each FISH spot and its associated pixels on every z-stack by particle analysis (ImageJ) ^40^. A FISH spot was called isolated from SC35 clusters only when none of its associated pixels exhibited SC35 signal. Otherwise a FISH spot was called co-localized with SC35. This is a conservative approach to call isolated FISH spots. Among the 210 nsaPeak FISH spots identified from 82 individual cells, 170 FISH spots (84.0%) co-localized with SC35 clusters. In comparison, among the 193 non-nsaPeak FISH spots identified from 100 cells, 111 co-localized with SC35, reflecting a 3-fold reduction in odds (odds ratio = 3.1, p-value < 5×10^−7^). We also summarized the proportion of co-localized FISH spots in each image. The 10 images stained with nsaPeak probe exhibited on average 81.1% of their FISH spots co-localized with SC35 (dots in left column, Figure 4D). In comparison, the 12 images (dots in right column) stained with the non-nsaPeak probe had on average 56.2% FISH spots co-localized with SC35 (p-value < 0.003, T test) (Figure 4D).

**Figure 4.**
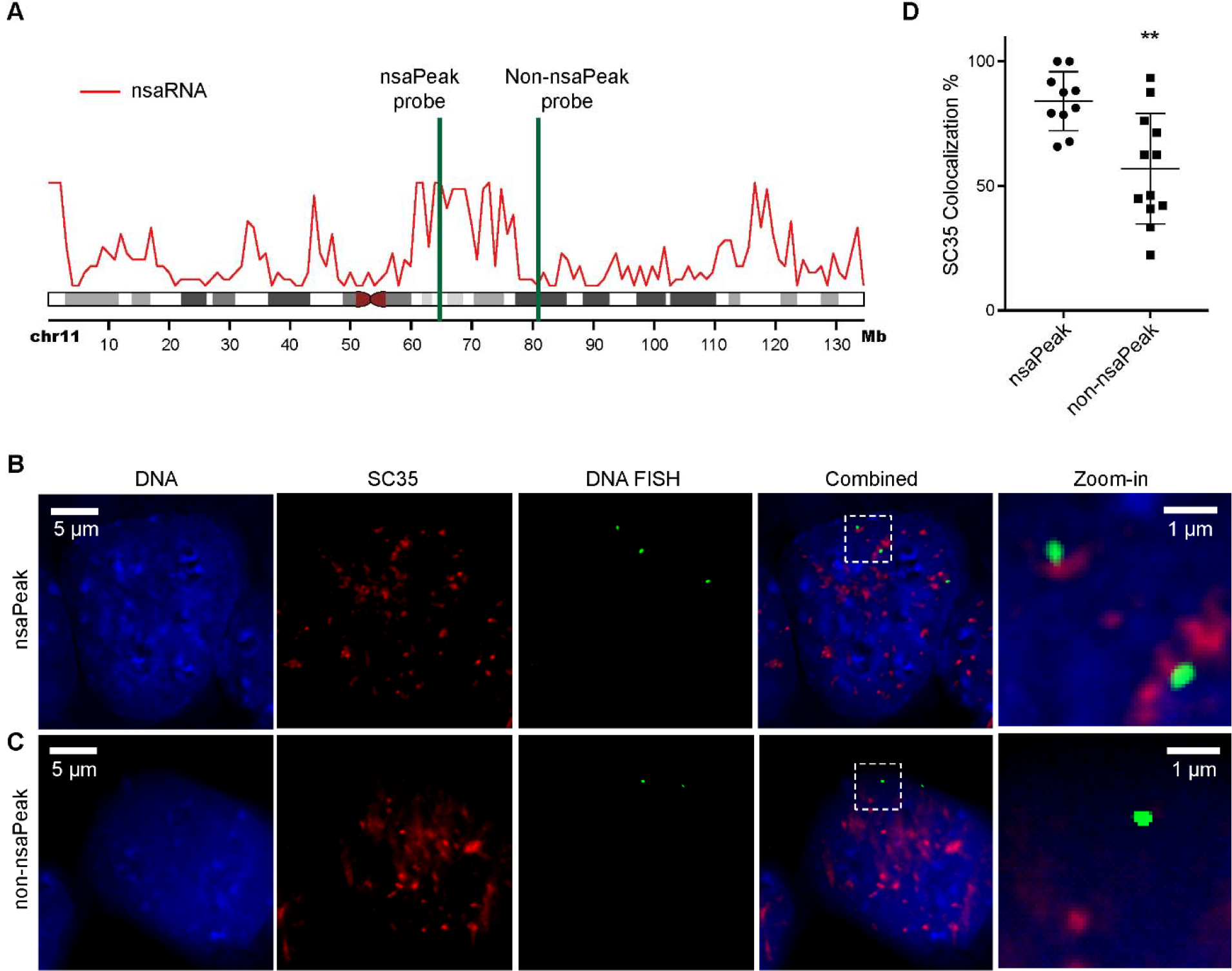
Visualization of representative nsaPeak and non-nsaPeak with SC35 clusters. (A) Genomic positions of nsaPeak probe and non-nsaPeak probe (green) with respect to nsaPeaks (red). (B-C) Representative images of HEK293T cells co-stained with Hoechst (DNA, blue), SC35 (red) and DNA FISH (green) with nsaPeak probe (B) and non-nsaPeak probe (C). Scale bar: 5 μm. Last column: zoom-in views of the selected regions in the dashed boxes. (D) The percentage of FISH spots that exhibited overlapping SC35 signal (y axis) in each image (dot) were plotted for samples interrogated with the nsaPeak probe images (left) and the non-nsaPeak probe (right). Error bar: standard deviation. **: p-value < 0.003.

In the second analysis, we compared the FISH-to-SC35 distance distributions between nsaPeak and non-nsaPeak samples. We computed center-to-center distance in 3D from every FISH spot to its nearest SC35 cluster. We summarized the number of center-pairs at each distance from 1 to 10 voxels in every image (Figure S6). The nsaPeak images exhibited 2 to 3 times more center pairs than non-nsaPeak images at every distance (p-value < 10^−5^, Kolmogorov test). For example, the nsaPeak images exhibited 1 to 18 center pairs at the distance of 8 voxels, whereas non-nsaPeak images exhibited 0 to 3 at this distance (Figure S6). The different distance distributions suggest that the interrogated nsaPeaks are closer to SC35 clusters than the interrogated non-nsaPeaks among the analyzed single cells. Taken together, the two analyses based on different analysis assumptions both revealed clear differences in relative positions of nuclear speckles to the two interrogated genomic regions. In summary, pre-mRNA data, CDK9 data, and single-cell image data supported the nsaPeaks as nuclear speckle proximal genomic regions.

### nsaPeaks are enriched in but not completely contained within A compartment

We exploited how nsaPeaks fit into current knowledge of 3D structure of the genome. Toward this goal, we compared nsaPeaks to nuclear compartments ^41^ and TADs ^42^. We called A/B compartments ^43^ from HEK293T Hi-C data ^44^ with Homer (v4.8.3) ^38,43^. Approximately half of the genome were associated with the A compartment (first row, Table S2). Approximately half of the genome in the A compartment and slightly more than 10% of the genome in the B compartment are associated with nsaPeaks (Figure 5), suggesting that nsaPeaks are enriched in but not a completely subset of the genomic sequences in the active compartment. In line with this observation, genes within nsaPeaks exhibited greater expression levels (Figure S7A). In addition, nsaPeaks exhibited baseline increase of H3K4me3, H3K27ac and H3K36me3 (25 – 75 quantiles, Figure S7B).

**Figure 5.**
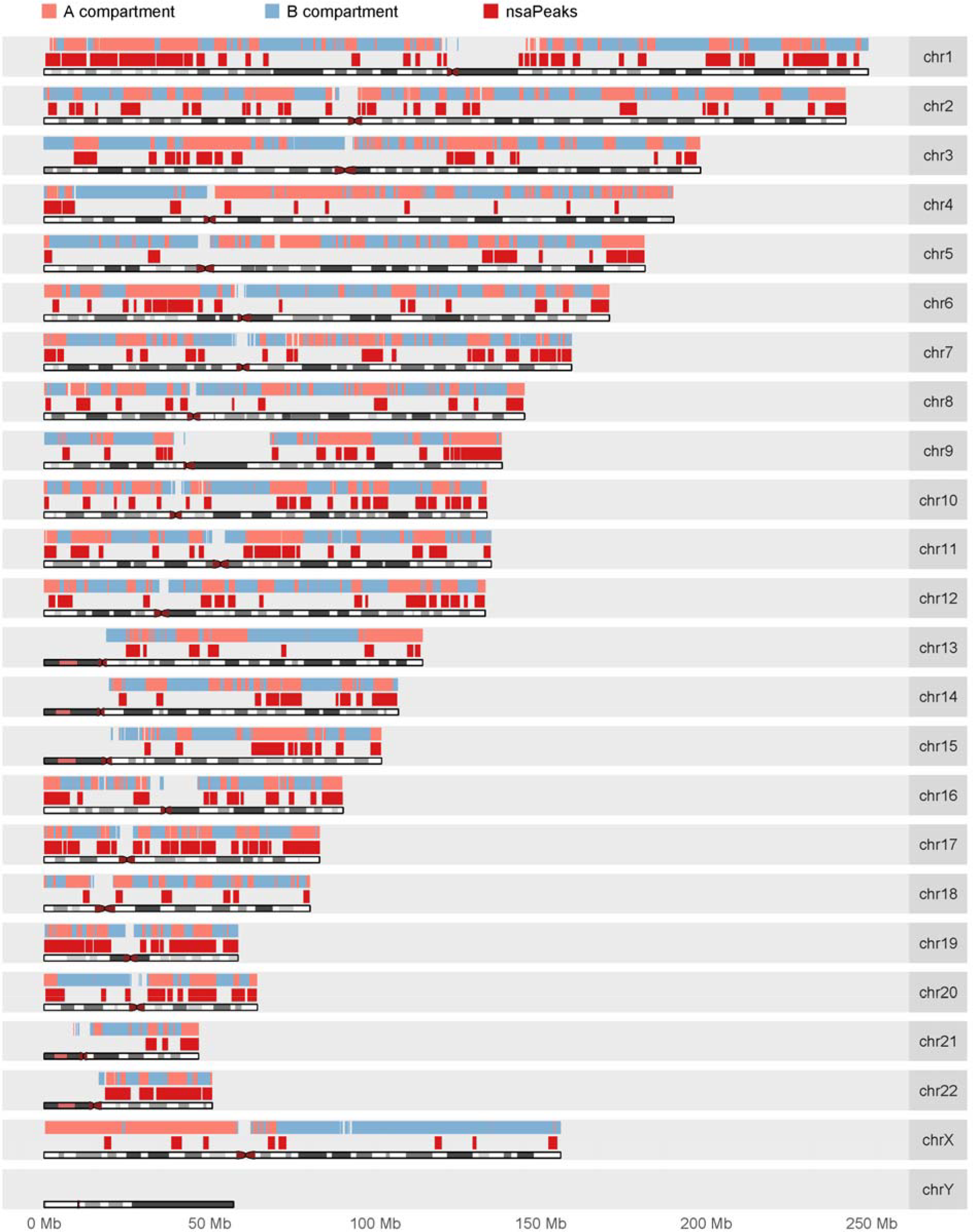
Genome-wide view of A (pink) / B (light blue) compartments and nsaPeaks (red).

### Genome sequence in a TAD tends to either entirely close to or distant from nuclear speckles

The notion of TADs was derived from Hi-C experiments ^42^ and TADs are subsequently proposed as a structural unit of genome organization ^45^. We reasoned that organizational units should exhibit unity in relative positions to other nuclear components, and therefore proximity of the genome and nuclear speckles may offer an alternative test to this proposition. We compared the 3,258 TADs derived from HEK293T Hi-C data (GEO: GSM1081530) ^44^ and nsaPeaks. Nearly 50% (289 out of 590) of the boundaries of nsaPeaks were aligned with TAD boundaries (p-value = 0.03, permutation test) (Figure 6A,B). Seventy-four nsaPeaks were aligned with 361 TADs, where each nsaPeak coincided with one TAD or several consecutive TADs (p-value = 0.051, permutation test). Recognizing the sensitivity of peak boundaries to noises in data and to algorithm, we did another test with an alternative set of boundaries. Based on the significant overlap of nsaPeaks and CDK9 broad peaks (Figure S5), we merged the two sets of peaks (union) and obtained 334 union-peaks. Approximately 52% (350 out of 668) of union-peak boundaries were aligned with TAD boundaries (p-value = 0.001, permutation test). Ninety-eight union-peaks were aligned with 468 TADs, where each union-peak coincided with one TAD or several consecutive TADs (p-value = 0.005, permutation test). Taken together, MARGI data suggest that the genomic sequence of a TAD tends to either entirely close to or entirely distant from nuclear speckles, supporting the proposition of TADs being structural units.

**Figure 6.**
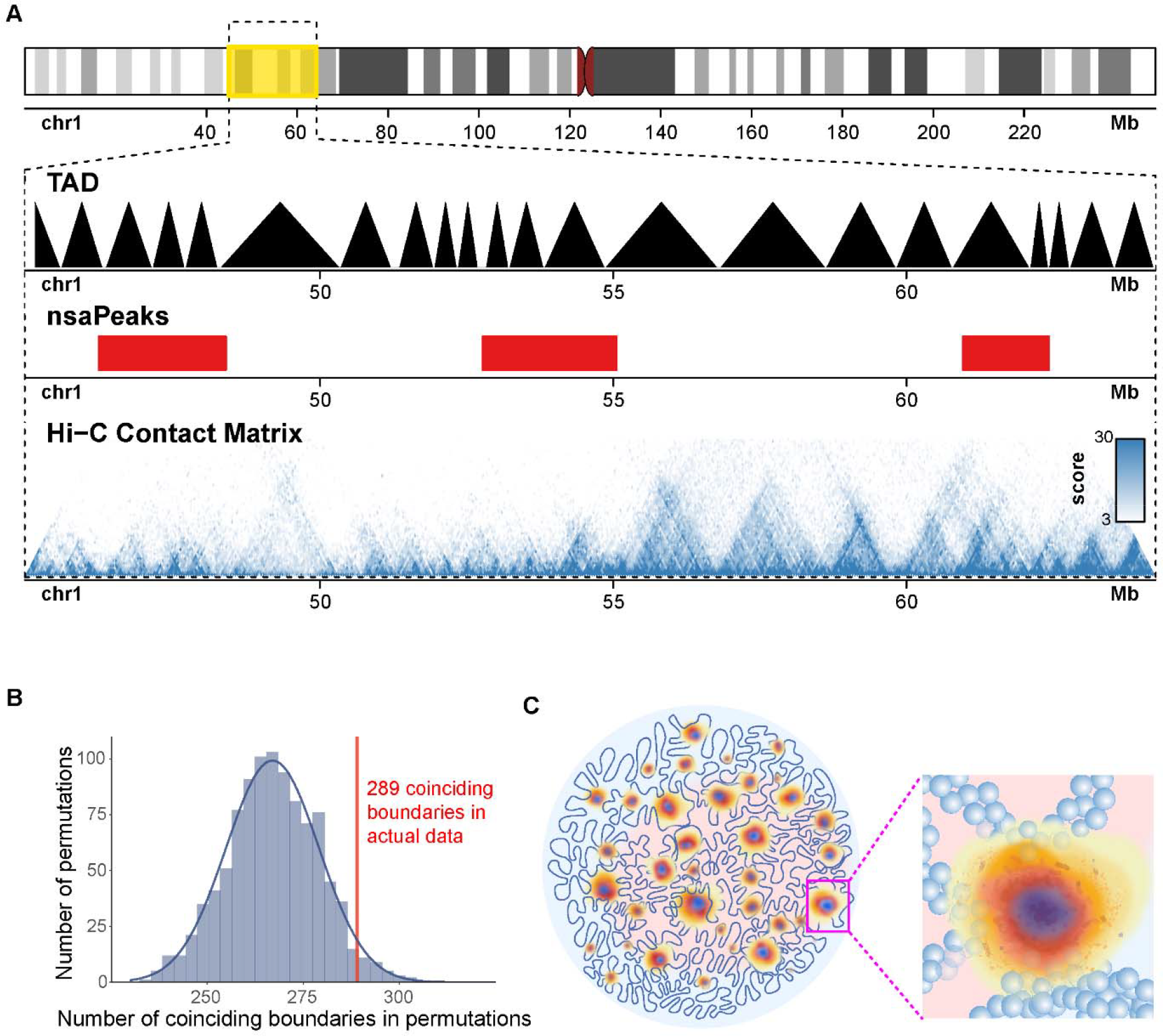
nsaPeaks and TADs. (A) Genome view of TADs, nsaPeaks, and Hi-C contact matrix. (B) Background distribution of the numbers of TAD boundaries coinciding with TAD boundaries from 1,000 permutations (histogram and fitted curve) versus the number of observed coinciding boundaries in actual data (red line). (C) A model of boundaryless nuclear speckles and the genome. Nuclear speckle cores are in red. Other nuclear speckle associated molecules exhibit diffusive patterns centered by nuclear speckle cores (blue, orange, yellow), and some of which extend to sufficient proximity to certain TADs (balls, insert). Pink/light blue: A/B compartment.

## Discussion

### Challenges in identifying relative positions of nuclear speckles with respect to genomic sequence

More than 150 proteins were reported to be associated with nuclear speckles ^46^, including snRNPs and SR proteins essential for RNA splicing ^47^ and a number of kinases and phosphatases that regulate splicing machinery ^7^. However, most of these proteins are not only present in nuclear speckles, and there is not sufficient data to assess the specificity of their localization to nuclear speckles. Therefore, the small number of proteins localized at the core of nuclear speckles, namely SC35 and SON received focal attentions and used as nuclear speckle markers in attempts to identify nuclear speckle-proximal genomic regions ^6–8^. However, the detachment of nuclear speckle cores to chromatin suggested that ChIP-seq analyses of nuclear speckle core proteins would unlikely reveal the genomic sequences close to nuclear speckles ^8^. Thus, finding relative positions of nuclear speckles with respect to genomic sequences remains a major challenge in nucleome research ^1^.

### RNAs as media for proximity labeling

The increasing evidence on “noncoding RNAs functioning as scaffolds in the construction of nuclear bodies” points to the essential role of RNA in nuclear bodies ^48^. Nuclear speckles exhibit clear centers but showed inconsistent boundary lines when visualized by staining different nuclear speckle markers ^9^. Evidence of nsaRNA locating at the periphery regions of nuclear speckles ^9^ fostered our hypothesis of this study that nsaRNAs serve as “proximity labeling” media, that “mark” proximal DNA (Figure 6C). Recently developed MARGI technology ^14^ enabled us to further examine this hypothesis by analyzing RNA-chromatin interactions of many noncoding RNAs at the same time.

### Cellular heterogeneity and assays of bulk cells

A rationale of ChIP-seq and ATAC-seq analyses of bulk cells is that if the majority of single cells share the same transcription factor binding regions or transposase accessible regions, and such commonality would be identified as peaks in bulk cell experiments. This rationale was verified by single-cell data produced by subsequently invented single-cell ChIP-seq ^49^ and single-cell ATAC-seq technologies ^50^. The same rationale is applicable to the MARGI technology in that only the genomic regions shared (relatively invariable) across many single cells would have a chance to appear in a bulk cell assay, whereas single cell-specific interaction regions can hardly produce significant signals in a bulk cell assay (Figure 1B). Although there does not exist a single-cell version of MARGI technology, single-cell imaging analysis provided data consistent with this rationale.

### Genome as a surrogate coordinate for studying nuclear organization

Revealing spatial organization of nuclear components has become a central task in nucleome research. This task is hindered by lack of a 3D coordinate system for the nucleus. Without a coordinate system, spatial data obtained from different single cells cannot be aligned, making it difficult to derive or test for any underlying principles.

Chromosome territories fill sizable portions of interphase nuclei ^24^. The correspondence between the any piece of uniquely mappable sequence and its genomic location makes it possible for the nuclear genome to serve as a surrogate coordinate system of the nucleus, given that a 3D location in the nucleus could be approximated by its nearest genomic sequence. Compared to the alternative of not having any 3D coordinate at all, the genome-surrogate-3D-coordinate provides a primitive means to record positional information which is potentially comparable across single cells or cell types. This surrogate coordinate has its own limitations, including lack of means to transform the surrogate coordinate into a physical coordinate and lack of power to differentiate chromosome pairs. This work was a test of this genome-surrogate-3D-coordinate. Both chromosomes and nuclear bodies could have variable and cell-specific 3D positions, however, our data suggested that the relative positions between nuclear speckles and chromosomes were relatively stable. Thus, accumulated knowledge of relative positions of various nuclear components ^34,51^ with respect to the nuclear genome may unleash the power of the genome-surrogate-3D-coordinate in future analyses of spatial organization of the nucleus.

## Materials and Methods

### Datasets and accession numbers

Public datasets used in this work are MARGI data from HEK293T cells (GEO: GSM2427902 and GSM2427903) and H9 hES cells (GEO: GSM2427895 and GSM2427896) ^14^, CDK9 ChIP-seq (GEO: GSM1249897) ^35^ control ChIP-seq (GEO: GSM2423406) ^37^ and Hi-C data from HEK293T cells (GEO: GSM1081530) ^44^, RNA-seq (GEO: GSM2155552) ^52^, H3K4me3 ChIP-seq (GEO: GSM945288, Encode: ENCFF001FJZ) and control ChIP-seq (GEO: GSM945256, Encode: ENCFF001HNC) from HEK293 cells ^53^, H3K4me1 ChIP-seq (Encode: ENCFF002AAV) ^54^, H3K9me3 ChIP-seq (Encode: ENCFF002AAZ) ^37^, H3K27ac ChIP-seq (Encode: ENCFF002ABA) ^54^, H3K36me3 ChIP-seq (Encode: ENCFF002ABD) ^37^, control ChIP-seq (GEO:GSM935586, Encode: ENCFF000WXY) ^37^ from HEK293 cells.

### Mapping MARGI data

After removing RCR duplicates, the RNA-end and the DNA-end of a read pair were separately mapped to the genome (hg38) using STAR (Version 2.5.1b) ^55^. Splice junction was allowed in mapping the RNA-end, by feeding the junction information (gtf file from ENSEMBL, hg38 release 84) to STAR. Splice junction was not allowed in mapping the DNA-end. Only the read pairs with both the RNA-end and the DNA-end uniquely mapped to the genome were used for downstream analysis.

### Identifying rRNA-DNA read pairs

Human rRNA genes include 45S (18S, 5.8S and 28S) in rDNA (human ribosomal DNA complete repeating unit, GenBank: U13369.1) as well as 5S and 5.8S in the human genome assembly (hg38) ^56^. A MARGI read pair is categorized as an rRNA-DNA pair when the RNA-end is uniquely mapped to any human rRNA gene and the DNA-end is uniquely mapped to a combined “genome” of hg38 concatenated with rDNA.

### Identifying nsaRNA-DNA read pairs

Human U1, U2, U4, U4atac, U5, U6, U6atac, U11, U12, 7SK, and Malat1 genes are considered nsaRNA genes. A MARGI read pair is categorized as an nsaRNA-DNA pair when the RNA-end is uniquely mapped to any human rRNA gene and the DNA-end is uniquely mapped to human genome (hg38). To minimize inclusion of nascent RNA, the read pairs with the RNA-end and DNA-end mapped to within 2,000 bp in the genome are removed from further analysis.

### Identifying pre-mRNA-DNA pairs

A MARGI read pair is categorized as a pre-mRNA-DNA pair when the RNA-end is uniquely mapped to an exon-intron junction with at least 10 bp overlap with the intron and the DNA-end is uniquely mapped to human genome (hg38). To minimize inclusion of nascent RNA, the read pairs with the RNA-end and DNA-end mapped to within 2,000 bp in the genome are removed from further analysis.

### Calling peaks and broad peaks

ChIP-seq and control ChIP-seq reads were mapped to human genome (hg38) and the uniquely mapped reads were fed to MACS2 ^36^ to call peaks. CDK9 broad peaks, pre-mRNA broad peaks, and nsaPeaks were identified by the findPeaks module in Homer (v4.8.3) ^38^. Any nsaPeak containing less than 9 MARGI reads was removed from further analysis.

### Calling TADs and A/B compartments

HEK293 Hi-C data (GEO: GSM1081530) ^44^ were aligned to hg38 retaining uniquely mapped reads. TADs were identified using a previously described HMM model ^42^ automated in the GITAR software ^57^. A/B compartments were called by the runHiCpca module in Homer (v4.8.3) ^38^.

### DNA FISH and immunofluorescence staining

The nsaPeak probe (RP11-772K10, covering chr11:64,663,168-64,947,112) with 5-ROX conjugate and the non-nsaPeak probe (RP11-908J16, covering chr11:80,767,575-80,980,051) with fluorescein conjugate were ordered from Empire Genomics. HEK293T cells were used through this study. In each experiment, cells were seeded on 18 X 18 mm glass coverslips with #1.5 thickness (#12-541A, Fisher Scientific) in 6-well tissue culture plate (Thermo Fisher Scientific) and grown in DMEM high-glucose media containing 10% (v/v) fetal bovine serum and 1% (v/v) penicillin-streptomycin at 37°C with ^5% CO^2^. Once reaches 80% confluency, the cells were rinsed with PBS and fixed with 4%^ paraformaldehyde (PFA) in pH 7.2 phosphate-buffered saline (PBS) for 30 min at room temperature. PFA was discarded and residual PFA was quenched by incubation with 0.1 M Tris buffer (pH 7.4) at room temperature for 10 min followed with one wash with PBS. Cells were permeablized with PBS containing 0.1% saponin (#84510-100, Sigma) and 0.1% TritonX-100 for 10 min, then with 20% glycerol in PBS for 20 min at room temperature with gentle shaking. Cells were rapidly frozen in liquid nitrogen and thawed at room temperature for three cycles, and rinsed with PBS. To detect SC35, cells were blocked with 5% bovine serum albumin (BSA) in PBS with 0.1% TritonX-100 (PBST) at 37°C for 30 min, and incubated with mouse monoclonal anti-SC35 antibody (1:250) (#Ab11826, Abcam) in blocking buffer at 37°C for 1 hour. Cells were washed with PBST for 10 min for twice with gentle shaking, incubated with goat anti-mouse IgG antibody conjugated with Alexa647 (1:200) (#A21236, Invitrogen) in blocking buffer at 37°C for 30 min, and then washed again with PBST for 10 minutes twice while shaking. The cells were fixed again with 2% PFA at room temperature for 10 min, quenched with 0.1 M Tris buffer as previously described and washed with PBS for 5 min. Cells were incubated with 0.1 M HCl for 30 min at room temperature, followed by 1 hr incubation with 3% BSA and 100 μg/mL RnaseA (#EN0531, Thermo Fisher Scientific) in PBS at 37°C. Cells were permeablized again with 0.5% saponin and 0.5% TritonX-100 in PBS for 30 min at room temperature with gentle shaking, and rinsed with PBS. Cells were further denatured by incubation in 70% formamide with 2X saline-sodium citrate (SSC) buffer at 73°C for 2 min 30 sec and then incubation in 50% formamide with 2X SSC at 73°C for 1 min. For each coverslip, 1.2 μL of FISH probes were mixed with 4.8 μL formamide, incubated at 55°C for 15 min and mixed with 6 μL 2X hybridization buffer (8X SSC with 40% dextran sulfate) followed with denaturation at 75 °C for at least 5 min until the cells were ready. 12 μL of FISH probe mixture was added onto a glass slide and quickly covered by freshly denatured coverslips with the cell side facing down. The coverslip was sealed with rubber cement and incubated in a humidified chamber at 37°C for 20 – 24 hr in the dark. The coverslips were collected the next day, washed twice with 50% formamide with 2X SSC for 15 min each at 37°C, three times with 2X SSC for 5 min each at 37°C, three times with 4X SSC containing 0.1% Tween20 for 5 min each at room temperature, with gentle shaking, and rinse with PBS. Cells were then stained with Hoescht 33342 (1:500) for 15 min followed with 5 min washing in PBS, mounted on slides with 80% glycerol in PBS and sealed with nail polish. Images in size of 1024 X 1024 were acquired on wide-field SIM DeltaVision Deconvolution Microscope using a 100X/1.40 oil objective (GE Healthcare Life Sciences) (pixel size = 0.66 μm). A series of z-stack images across the cells were acquired with thickness of 0.15 μm. Deconvolution was performed on these Z-stacks for subsequent image analysis.

### Co-localization analysis

Deconvoluted images for each field of view contain a series of z stacks in three channels, DAPI, FISH and SC35. FISH spots were identified by performing particle analysis on the 2D maximal projection of z stacks of each field of view in the FISH channel with the threshold being set to the minimal value allowing only FISH spots to be recognized as “particles” in the size range of 10 – 500 pixels. These FISH particle regions were saved and applied to the z-stacks of FISH and SC35 channels and z-axis profiles of selected regions (min, max and mean values of fluorescence intensity) in both channels were recorded and examined. Positive co-localization of a given FISH spot with SC35 was defined by the presence of positive SC35 signals above background in any of the FISH-signal containing regions of that FISH spot. In order to determine one FISH region as positive SC35-colocalized, it needs to contain more than one stack with mean intensity above SC35 background, or contain more than half amount of stacks with max intensity over SC35 background. For each analyzed image, SC35 background value was based on the average mean intensities in areas outside of SC35 clusters within the nucleus region. For each field of view, the SC35 co-localization rate represents the ratio of the amount of SC35-colocalized FISH spots over the amount of total FISH spots.

Center-to-center distances were calculated as follows. After deconvolution, each cluster or spot was identified as a connected 3D region such that all voxels within this region are above a threshold. The threshold was determined as described previously ^58^. Briefly, each deconvoluted image was scanned to identify all pixels on every stack that was could not possibly be background. The threshold was chosen such that the number of detected fluorescent clusters would not change within 3% variation of this threshold. The center of a cluster (spot) was calculated as the gravity center. Center-to-center distance was calculated with voxel as the unit.

## Acknowledgment

Authors thank Dr. Alexandra Bortnick for advices on optimizing the immunofluorescence and DNA FISH assay. This work is funded by NIH R35CA197622 (J.Z.), DP1HD087990 (S.Z.) and U01CA200147 (S.Z.).

